# INFLUENCE OF A 30-DAY SPACEFLIGHT ON THE STRUCTURE OF MOTONEURONS OF THE TROCHLEAR NERVE NUCLEUS IN MICE

**DOI:** 10.1101/2020.09.16.299461

**Authors:** Lyubov Pavlik, Irina Mikheeva, Rashid Shtanchaev, Gulnara Mikhailova, Vladimir Arkhipov

## Abstract

During spaceflight and immediately after it, adaptive neuroplastic changes occur in the sensorimotor structures of the central nervous system, which are associated with changes of mainly vestibular and visual signals. It is known that the movement of the eyeball in the vertical direction is carried out by muscles that are innervated by the trochlear nerve (CN IV) and the oculomotor nerve (CN III). To elucidate the cellular processes underlying the atypical vertical nystagmus that occurs under microgravity conditions, it seems necessary to study the state of these nuclei in animals in more detail after prolonged space flights. In this work, we carried out a qualitative and quantitative light-optical and ultrastructural analysis of the nuclei of the trochlear nerve in mice after a 30-day flight on the Bion-M1 biosatellite and followed by a stay for 13-14 hours under the influence of the Earth’s gravity after landing. As a result, it was shown that the motoneurons in the nucleus of the trochlear nerve changed their morphology under the influence of microgravity. Cell nuclei of the motoneurons had a more simplified rounded shape than in the control. In addition, the dendrites of these motoneurons significantly reorganized geometry and orientation under microgravity conditions; the number of dendritic branches has been increased to enhance the reduced signal flow. Apparently, to ensure such plastic changes, the number and size of mitochondria in the soma of motoneurons and in axons coming from the vestibular structures increased. In addition, the experimental animals showed an increase in the size of the cisterns of the rough and smooth endoplasmic reticulum in comparison with the control group of animals left on Earth, for which the environmental conditions in the spacecraft were reproduced. Thus, the main role in the adaptation of the trochlear nucleus to microgravity conditions, apparently, belongs to the dendrites of motoneurons, which rearrange their structure and function to enhance the flow of sensory information. These results are useful for the development of new, more effective means to facilitate the stay and work of space travelers in a long spaceflight.

## Introduction

During a long-term spaceflight, a significant change in vestibular function occurs, which can lead to the development of space adaptation syndrome and space motion sickness [1–3]. These changes occur in space travelers both during spaceflights and during their return to Earth, but the cellular brain organization of such adaptation is still poorly understood. One of the manifestations of space motion sickness includes disorders of oculomotor reactions: increased oculomotor activity of a saccadic and smooth nature; inhibition of the tracking function of the eyes and the appearance of additional saccadic movements during smooth tracking with a transition to nystagmus-like reactions. In a person who has been in a long-term spaceflight, spontaneous atypical nystagmus (especially vertical) and a decrease in the thresholds of optokinetic nystagmus are also observed [3]. It has been shown that most disorders of oculomotor reactions are reversible, for example, atypical vertical nystagmus disappears 5-8 days after landing [4]. It is known that muscles, which are innervated by the trochlear nerve (CN IV) and the oculomotor nerve (CN III), carry out the movement of the eyeball in the vertical direction. The motoneurons of the nucleus of the trochlear nerve move the eyeball upward during vertical saccadic movements by innervation of the superior oblique muscle [5]. To elucidate the cellular processes underlying the atypical vertical nystagmus that occurs under microgravity, it seems necessary to study in more detail the state of these motoneurons in animals after prolonged spaceflights. These results will be useful for the development of new, more effective means to facilitate the stay and work of space travelers in a long-term spaceflight. We have previously carried out studies of motoneurons and neuropile of the oculomotor nucleus of the mouse brain [6]. In the present work, a morphometric study of the nucleus of the trochlear nerve was carried out at the light-optical and ultrastructural levels after a 30-day orbital flight of mice on a biosatellite in the framework of the Bion-M1 program.

## Results

### Histological investigations

Light-optical study of the motoneurons of the trochlear nerve nucleus on preparations obtained from animals after spaceflight did not reveal any noticeable changes in the structure of the motoneurons of the nucleus compared to the control group. In animals of both groups, the sizes of the soma diameter, nucleus, as well as the ratio: nucleus area/cytoplasm area did not differ significantly (S1Table). However, the most noticeable changes affected the geometric parameters of the dendrites. Three-dimensional reconstruction showed that the length of the proximal area of dendrites (before bifurcation) in the animals of the experimental group decreased compared to the control, and the number of branches in them was significantly greater (Fig.1). Moreover, the reorganization of the direction of motoneuronal dendrites occurs in such a way that the number of dendrites in the dorsal direction is greater, and in the medial direction it is less in experimental animals than in control animals (S1 Table).

**Fig 1.**
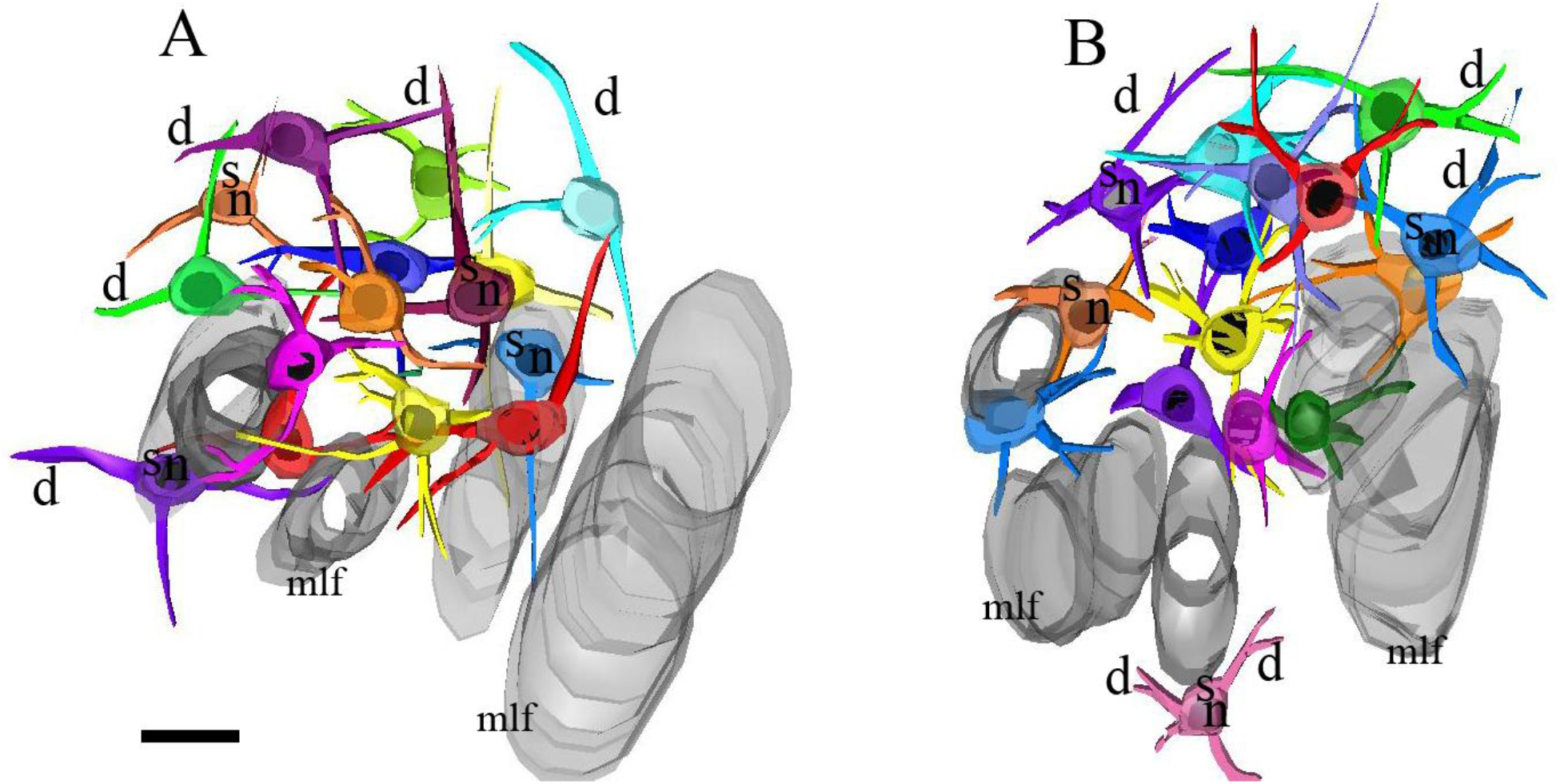
Three-dimensional reconstruction of motorneurons of the trochlear nucleus in the control (A) and experimental (B) groups. s-soma, n-nucleus, d-dendrites, mlf-medial longitudinal fasciculus. It can be seen that the dendrites of the motoneurons of the nucleus in experimental animals have more branches. Bar 50 μm

### Electron microscopy

As a result of ultrastructural studies, it was shown that morphological changes in motoneurons of trochlear nuclei in experimental animals are primarily related to biosynthetic processes. The number of mitochondria in the soma has increased, and their perimeters in the images have increased markedly (S2 Table). The size of the cisterns of the rough and smooth reticulum (Fig. 2B) was larger than in the control (Fig.2A), on average by 32 ± 7% and 33 ± 6%, respectively (S2 Table), which indicates an increase in the biosynthetic activity of the neuron.

**Fig 2.**
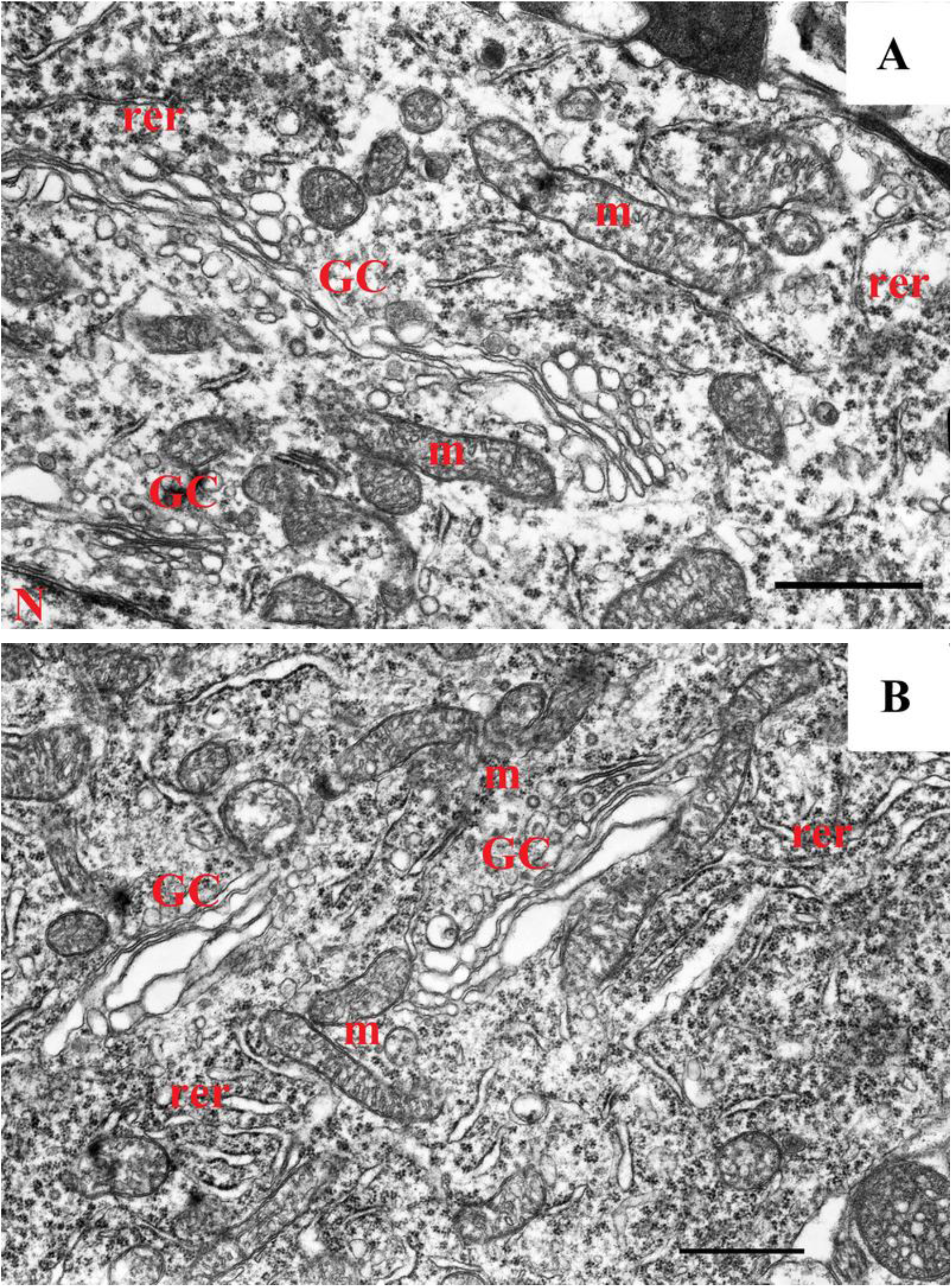
Ultrastructure of the soma of the trochlear nucleus motoneurons in the control (A) and in the experimental (B) groups. In animals of the experimental group, the rough reticulum has a more complex morphology than in the control group. In addition, there are more mitochondria and their size is larger. N – nucleus, m – mitochondria, rer – rough endoplasmic retiuculum, GC – Golgi complex. Bar 1 μm

The neuropil of the trochlear nerve nucleus in mice, like in other animal species [7] consists of axo-somatic synapses and several types of axo-dendritic synapses. The neuropil contains axo-dendritic synapses with vesicles that are very different in size, as well as synapses with vesicles of approximately the same size, 35–40 nm, with the inclusion of dense core vesicles (Fig. 3). The study of the ultrastructure of axodendritic synapses showed that in control animals (Fig. 3A) the vesicles in the synaptic terminal are evenly distributed, while in experimental animals a significant part of the vesicles are grouped near the active zone (Fig. 3B).

**Fig 3.**
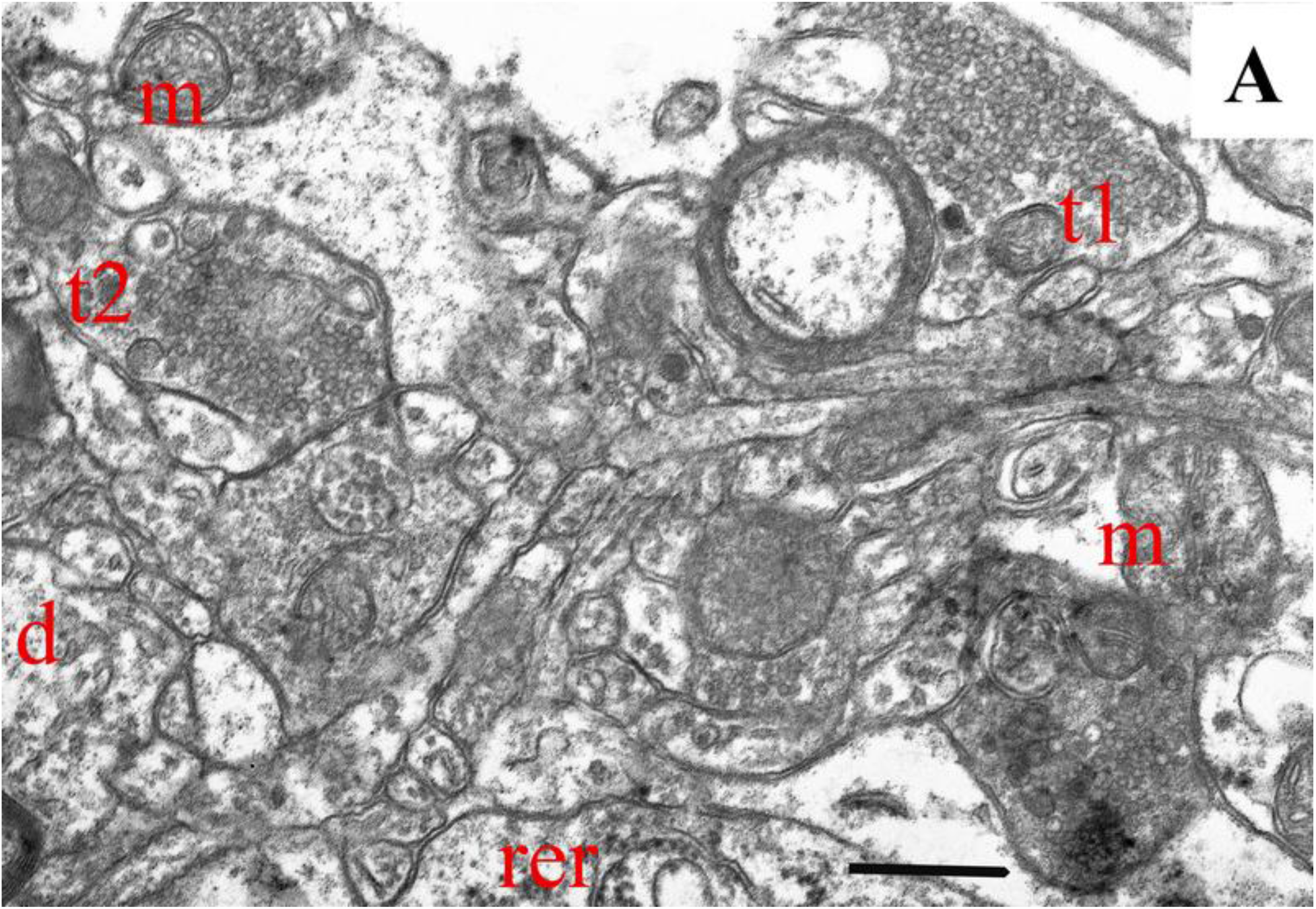
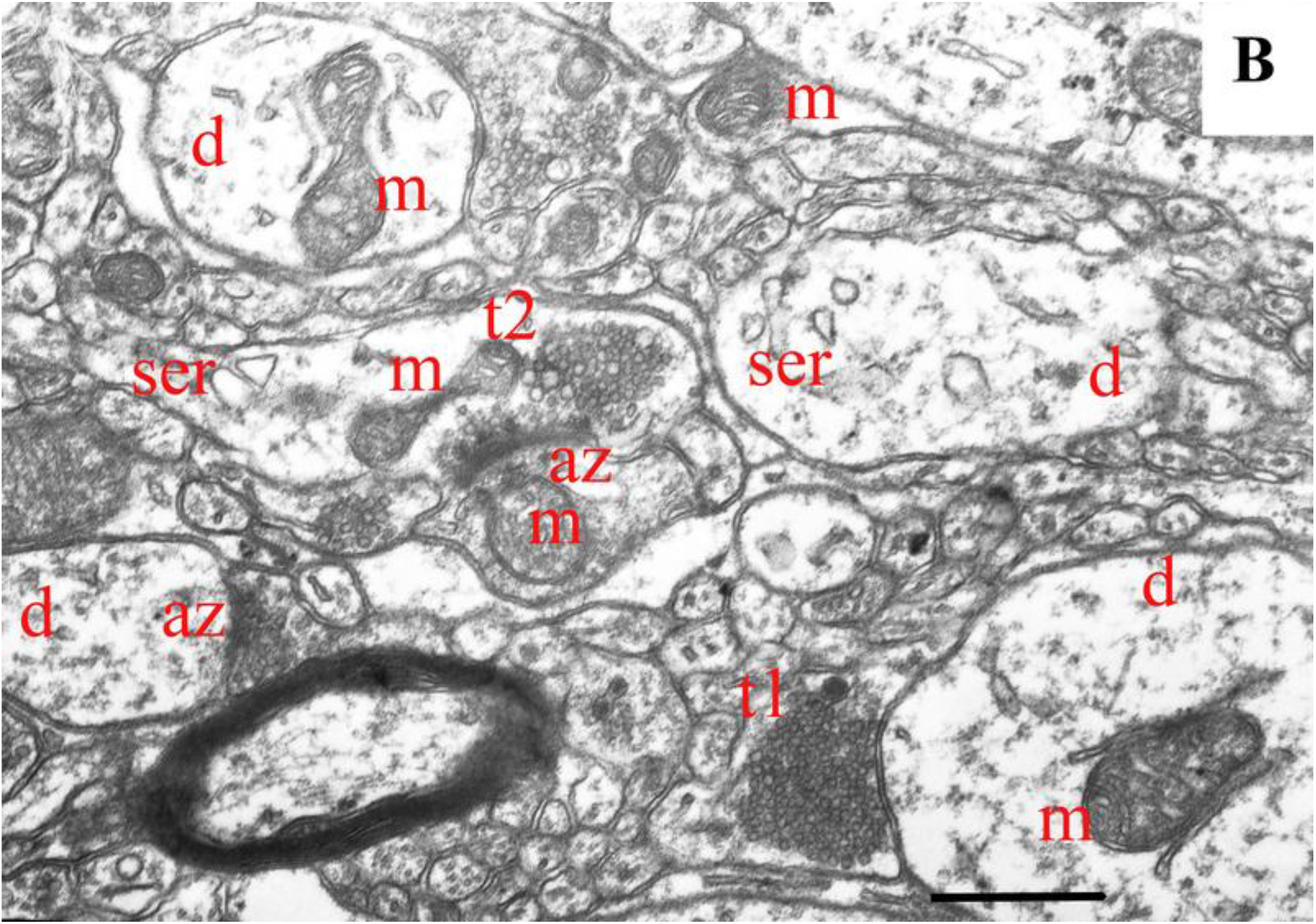
Ultrastructure of the neuropil of the trochlear nucleus motoneurons in the control (A) and in the experimental groups (B). In animals of the control and experimental groups, one can see two types of presynaptic endings of axodendritic synapses (t1 and t2), differing in size and distribution of vesicles: presynaptic endings of t1 have vesicles 35-40 nm with the inclusion of dense-core vesicles; t2 endings additionally have large vesicles up to 100 nm. It can be seen from the images that in animals of the experimental group (B), the distribution of vesicles in the presynaptic terminal is uneven, with grouping near the active zone, in contrast to motoneurons in control animals (A), the distribution of vesicles in which is uniform. Mitochondria in the trochlear nucleus of experimental animals (B) are more numerous and among them there are a large number of dividing mitochondria. az – active zone, d – dendrite, ser – smooth endoplasmic reticulum, m – mitochondria, Bar 1 μm

Mitochondria in the dendrites of motoneurons, as in the soma in experimental animals, have retained their integrity. Moreover, their number and size were significantly larger than that of the control animals (S2 Table).

## Discussion

In this work, we studied the morphology of motoneurons and neuropil of the trochlear nerve nucleus (CN IV) of mice after a 30-day flight on the Bion-M1 biosatellite, followed by a stay of 13-14 hours under the influence of the Earth’s gravity after landing. The flight significantly influenced the behavior of animals: approximately two hours after landing, the animals showed signs of impaired motor function [8]. The mice could not maintain a normal posture, their physical activity was significantly reduced; when tested in a drop test, they were not able to roll over. The animals had a pronounced exophthalmos, which persisted for a long time, in contrast to the motor activity, which was restored 6-8 hours after landing. One of the reasons for visual-vestibular disorders identified after spaceflight in humans and animals (specifically, atypical nystagmus in humans) is functional disorders of the oculomotor nucleus and nucleus of the trochlear nerve caused by vestibular deprivation. Our previous results showed that spaceflight caused a number of morphological changes in the oculomotor nucleus, which mainly concerned the structure of axo-dendritic synapses [6]. A decrease in the number of axo-dendritic synapses was compensated by an increase in the number of active synapses, which followed from an increase in the length and width of the postsynaptic density [6].

In present work, we investigated the ultrastructural features of the motoneurons and neuropil of the trochlear nucleus, which also be involved in the mechanisms of disturbances caused by spaceflight. The results obtained revealed adaptive changes in this structure of the animal brain. Although the sizes of the soma and cell nucleus of motoneurons in animals of the experimental and control groups did not differ, nevertheless, the morphology of the soma and dendrites differed significantly (S1 Table, S2 Table). In animals of the experimental group, morphological changes occurred in the endoplasmic reticulum and mitochondria. In the trochlear nerve nucleus of animals in the experimental group cistern perimeters of the rough and smooth reticulum increased significantly. This indicates an intensification of biosynthetic processes in motoneurons, since posttranslational modification of protein molecules and lipids biosynthesis occurs in the endoplasmic reticulum [9]. In addition, one of the main functions of the endoplasmic reticulum is the absorption and deposition of cytoplasmic calcium entering the cell [9, 10]. Given that the experimental animals were under the influence of gravity for 13 hours after space flight, it can be assumed that this also contributes to the adaptive rearrangement of the endoplasmic reticulum. Interestingly, a significant increase and complication of the cisterns of the endoplasmic reticulum, as well as an increase in the size of mitochondria, were noted by other researchers in Purkinje cells of rats after 24 hours of space flight [11]. Mitochondria, along with the reticulum, also serve as a Ca^2+^ store in the cell [12]. An increase in the perimeter of mitochondria and their number in the soma of neurons and in the axonal endings in the neuropil of the trochlear nucleus indicates an increased ability of mitochondria to bind excess Ca2 ^+^, which is an additional adaptation to microgravity. Mitochondria are the most labile intracellular structures [13,14]. They are among the first to change with increased activity of neurons and with damaging influences [15–20].

When investigating the neuropil of the trochlear nerve nucleus in experimental mice, we found dendrites altered to varying degrees. It is obvious that the state of these dendrites reflects a long-term adaptation to the absence of vestibular afferentation during flight. In our work, this was expressed in the appearance of additional branches of dendrites and a change in their direction (Fig.1). The revealed changes are consistent with the ultrastructural data of other authors, who revealed a decrease in the functional activity of afferent inputs in microgravity in brain structures [1, 21, 22]. For example, similar result was obtained in the somatosensory cortex of the rat brain under microgravity conditions on the 12th day of flight [23]. We believe that the described changes are reversible, since oculomotor disturbances usually disappear some time after landing [3]. It is known that natural deprivation of visual and vestibular inputs causes a change in the morphology of dendrites receiving signals from sensory inputs [24]. Thus, the revealed morphological changes in the motoneurons and neuropil of the nucleus of the trochlear nerve contribute to the mechanisms of oculomotor disorders caused by prolonged spaceflight.

### Conclusions

Under microgravity conditions, there is a significant decrease in the inflow of information from the vestibular system to the motoneurons of the oculomotor nucleus and trochlear nucleus. Adaptive changes in the structural and functional properties of these motoneurons occur in such a way that the efficiency synaptic transmission increases. We revealed such changes in relation to the nucleus of the trochlear nerve and the oculomotor nucleus. In the oculomotor nucleus, this efficiency is mainly due to changes in the morphology of synapses [6], and in the nucleus of the trochlear nerve, due to changes in the structure of dendrites, which is the main result of this study. Dendrites are the basis for the plasticity of the mature brain under various influences, and therefore it is not surprising that they react markedly to microgravity. The results of the study of motoneurons of the trochlear nucleus in the present work showed that their dendrites did indeed undergo a number of changes. The increase in branching and orientation of dendrites that we found contributes to the adaptive enhancement of reduced signaling. The activation of the synthesis of proteins and lipids and the restructuring of the mitochondrial apparatus promote adaptive changes in motoneurons in response to microgravity.

## Materials and methods

### Bioethics

The bioethical examination of the study Protocol was conducted By the Commission on bioethics of the MSU mitoengineering research Institute (Protocol No. 35 of November 1, 2012) and the Commission on biomedical ethics of the SSC RF – IBP RAS (Protocol No. 319 of April 4, 2013). The experiments were carried out in accordance with the European Convention for the protection of vertebrates used for experiments or other scientific purposes (Strasbourg, March 18, 1986) [25] and order No. 742 of the Ministry of higher and secondary special education of the USSR “on approval of the Rules for carrying out work using experimental animals” dated 13.11.1984.

### Animals

Mice C57/BL6 became the main object of biomedical research in the Bion-M1 program, which have a number of advantages as a space biology object. The C57/BL6 mouse line (N in the full name of the line stands for the US national institutes of health subline) is one of the most widely used and was selected based on the wishes of the project participants. Male C57/BL6N mice numbering 300 individuals were obtained from the branch of the Institute of Bioorganic chemistry academicians M. M. Shemyakin and Yu. A. Ovchinnikov - nursery of laboratory animals “Pushchino” at the age of 8-9 weeks.

The experimental group consisted of mice weighing 29.3 ± 1.1 g; the age of the animals at the time of the launch of the biosatellite was 19 weeks.

The experimental group of mice (n=5) was on board the Bion-M 1 biosatellite, the control group of animals (n=9) remained on Earth, for which the environmental conditions in the spacecraft were reproduced .In 13-14 hours after landing, mice of the experimental group were euthanized by the method of cervical dislocation chosen as the least contrary to the implementation of the tasks of scientific Programs. [26] The brain was removed and a fragment containing the nucleus of the trochlear nerve was isolated.

### Histological investigations

The isolated brain fragment was placed in a fixing solution containing 2.5% glutaraldehyde in 0.1 M phosphate buffer. For histological and ultrastructural study of the tissue, we used methods with our modifications, which were described earlier [6]. Fragments of the brain were embedded in eponic resin, and frontal serial histological slices (5 μm) were prepared from the obtained blocks on a microtome. The trochlear nerve nucleus was identified on histological sections using stereotaxic atlases of the mouse brain [27] (Fig. 4).

**Fig 4.**
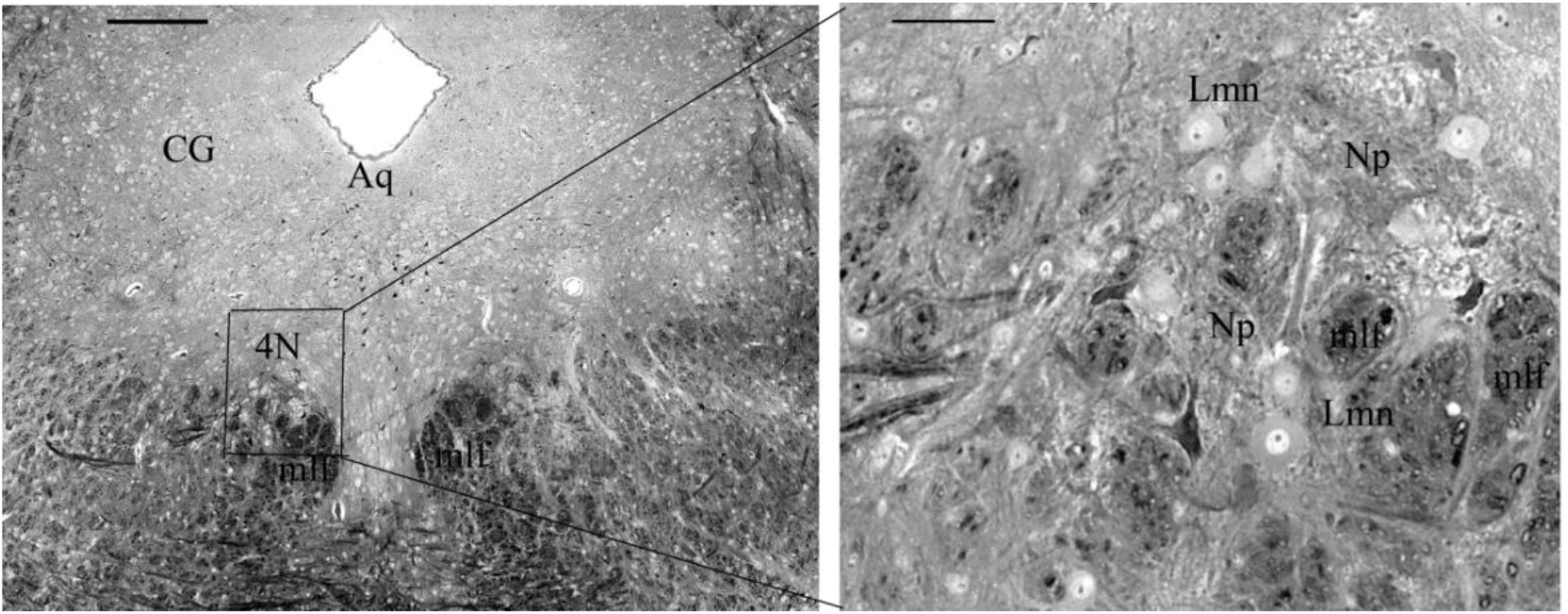
Coronary histological slice (5 μm) of the mouse brain containing the trochlear nucleus. On the left is the trolear nucleus, on the right it is enlarged. Aq – aqueduct, CG – central gray, 4N – trochlear nucleus, Lmn – large motoneurons, Np – neuropil, mlf – medial longitudinal fasciculus. Bar 300μm, insert-60μm

For further histological examination, we selected slices containing large motoneurons (with a soma diameter of about 20-25 μm), with clearly distinguishable boundaries of cells, nuclei, and nucleoli. Each 5 μm slice contained, as a rule, 1-6 central neurons, including the soma, nucleus, and nucleolus on each side of the brain. Neurons with a soma diameter less than 10-15 μm were not taken into account.

### Three-dimensional reconstruction of motoneurons

Three-dimensional reconstruction of the motoneurons and reference structures was performed using serial histological slices according to the method previously described [28–30]. The diameters of the soma and nucleus were determined on panoramic images of histological slices using Adobe Photoshop and PTGui v.9.1.8 Pro (USA). The number and direction of dendrites were evaluated. To determine the direction of the main dendrites, neurons with visible initial segments of dendrites were outlined in the images obtained on successive 5-6 slices with a thickness of 3-5 μm.

### Electron microscopy

Histological slices selected for further electron microscopy were glued onto a new epon block, the area with large motoneurons was targeted, and ultrathin sections (70-75nm) were obtained using a Leica EM UC6 microtome (Germany). Sections were contrasted with uranyl acetate and lead citrate and studied using a JEM-100B electron microscope (Japan). Ultrastructural morphometric analysis was carried out on photonegatives using the Image Tool (2013, England). In the soma of neurons, the perimeter of the cisterns of the rough endoplasmic reticulum and the smooth endoplasmic reticulum, the perimeter of the mitochondrial profiles were determined. In the neuropil, the number of axo-dendritic synapses with asymmetric postsynaptic density was assessed, as well as the number of active zones in synapses. In addition, the length of individual active zones, and the number and perimeter of mitochondrial profiles at the terminals of axo-dendritic synapses were assessed.

### Statistical analysis

The quantitative processing of the obtained results was carried out using Graph Pad for Prizma 5 and SigmaPlot 11.0 (Systat Software, Inc., 2008). The Student’s t-test was used to compare the experimental and control groups. In some cases (when data failed the equal variance test of Brown-Forsythe), we estimated the significance of differences using nonparametric single-factor dispersion analysis for repeated measurements (Kruskal–Wallis One Way Analysis of Variance on Ranks) which was followed with the pairwise comparison by the Tukey’s test. Quantitative results are presented in the text and tables as mean ± standard error of the mean.

## Supporting information

**S1 Table.** Morphometric analysis of motoneurons of the trochlear nucleus in the control and experimental groups

| Structure | Control group |  | Experimental group |  |
| --- | --- | --- | --- | --- |
| | Volume, $\times 10^3 \mu\text{m}^3$ | | | |
| soma | 16530±1448 | | 10317±1664 | |
| nucleus | 2450±49,7 | | 2218±77* | |
| Number of dendrites<br>(before bifurcation ) | 56 |  | 54 |  |
| number of branches | 45 |  | 71* |  |
|  | % of the total | Dendrite length<br>(before bifurcation) | % of the total | Dendrite length<br>(before bifurcation) |
| Lateral dendrites | 30,4 | 43,9±3,3 | 29,6 | 26,3±2,6** |
| Medial dendrites | 23 | 40,9±3,4 | 12,9 | 28,5±3,9* |
| Dorsal dendrites | 17 | 46,5±5,7 | 28 | 32,6±2,3* |
| Ventral dendrites | 28,5 | 56,4±2,7 | 29,6 | 32,4±1,7** |

\* The differences were considered statistically significant at $P < 0,05$ .
\*\*The differences were considered statistically significant at $P < 0,001$ .

**S2 Table.**
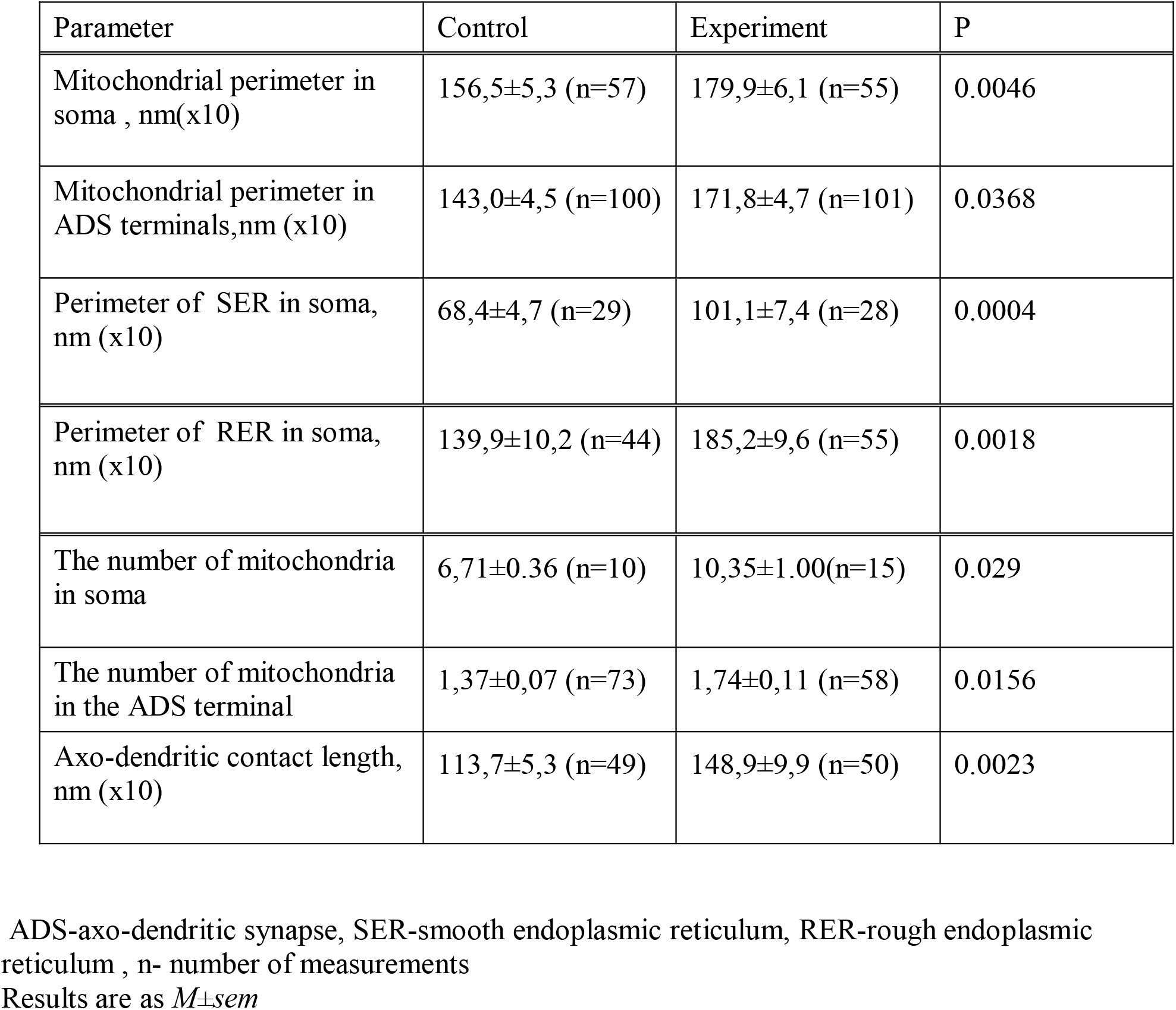
Morphometric analysis of intracellular organelles and neuropil of the mice trochlear nucleus in the control and experimental groups.

## Acknowledgment

This work was supported by the State task of ITEB RAS No. 075-00487-19-01

## References

1. Krasnov IB. Gravitational neuromorphology. Review. Adv Space Biol Med. 1994; (4):85–110. PMID: 7757255

2. Kornilova LN, Kozlovskaya IB. Neurosensory mechanisms of space adaptation syndrome. Human Phisiology. 2003; 29(5):527–538. https://doi.org/10.1023/A:1025899413655

3. Kornilova LN, Naumov IA, Glukhikh DO, Ekimovskiy GA, Pavlova AS, Khabarova VV, Smirnov YI., Yarmanova EN. Vestibular function and space motion sickness. Human Physiology. 2017; 43(5):557–568. https://doi.org/10.1134/S0362119717050085

4. Kornilova LN, Alekhina MI, Temnikova VV, Sagalovich SV, Malakhov SV, Naumov IA, Kozlovskaya IB, Reshke M, Vasin AV. The effect of a long stay under microgravity on the vestibular function and tracking eye movements. Human Physiology. 2006; 32(5): 547–555. https://doi.org/10.1134/S0362119706050082

5. Büttner-Ennever J. A. The extraocular motor nuclei: organization and functional neuroanatomy. Prog Brain Res. 2006; 151: 95–125. PMID: 16221587 DOI: 10.1016/S0079-6123(05)51004-5

6. Mikheeva IB, Shtanchaev RSh, Pen’kova NA, Pavlik LL. Structure of Interneuronal Contacts in the Neuropil of the Oculomotor Nuclei in Mouse Brain under Conditions of Long-Term Microgravity. Bull Exp Biol Med. 2018;165 (4):457–460. PMID: 30121909 DOI: 10.1007/s10517-018-4193-8

7. Bak IJ, Baker R, Choi WB, Precht W. Electron microscopic investigation of the vestibular projection to the cat trochlear nuclei. Neuroscience. 1976; 1(6):477–82.PMID: 11370240 DOI: 10.1016/0306-4522(76)90099-3

8. Andreev-Andrievskiy A, Popova A, Boyle R, Alberts J, Shenkman B, Vinogradova O, Dolgov O, Anokhin K, Tsvirkun D, Soldatov P, Nemirovskaya T, Ilyin E, Sychev V. Mice in Bio-M1 space mission: training and selection. PLoS One. 2014; 9(8):e10483 PMID: 25133741 PMCID: PMC4136787 DOI: 10.1371/journal.pone.0104830

9. Kucharz K, Wieloch T, Toresson H. Fission and fusion of the neuronal endoplasmic reticulum. Translational Stroke Research. 2013; 4 (6): 652–662. PMID: 24323419 DOI: 10.1007/s12975-013-0279-9

10. Karagas Nicholas E, Venkatachalam Kartik. Roles for the Endoplasmic Reticulum in Regulation of Neuronal Calcium Homeostasis. Cells. 2019; 8(10):1232. PMID: 31658749 PMCID: PMC6829861 DOI: 10.3390/cells8101232

11. Holstein G.R., Kukielka E., MartinelliG.P.Anatomical observations of the rat cerebellar nodulus after 24 hr of spaceflight. J Gravit Physiol. 1999; 6 (1): 47–50.PMID: 11543023

12. Cali Tito, Ottolini Denis, Brini Marisa. Mitochondrial Ca(2+) and neurodegeneration. Cell Calcium. Review. 2012; 52(1):73–85. PMID: 22608276 PMCID: PMC3396847 DOI: 10.1016/j.ceca.2012.04.015

13. Westermann Benedikt. Bioenergetic role of mitochondrial fusion and fission. Biochim Biophys Acta. 2012; 1817(10):1833–1838. PMID: 22409868 DOI: 10.1016/j.bbabio.2012.02.033

14. Mitra Kasturi. Mitochondrial fission-fusion as an emerging key regulator of cell proliferation and differentiation. Bioessays. 2013; 35(11):955–64.PMID: 23943303 DOI: 10.1002/bies.201300011

15. Di Lisa F, Canton M, Menabò R, Kaludercic N, Bernardi P. Mitochondria and cardioprotection. Heart Fail Rev. 2007; 12 (3-4):249–260.PMID: 17516167 DOI: 10.1007/s10741-007-9028-z

16. DuBoff B, Feany M, Götz J. Why size matters - balancing mitochondrial dynamics in Alzheimer’s disease. Trends Neurosci. 2013; 36(6):325–335. PMID: 23582339 DOI: 10.1016/j.tins.2013.03.002

17. Bernard-Marissal N, Chrast R, Schneider BL. Endoplasmic reticulum and mitochondria in diseases of motor and sensory neurons: a broken relationship? Cell Death Dis. 2018; 9 (3):333. PMID: 29491369 PMCID: PMC5832431 DOI: 10.1038/s41419-017-0125-1

18. Ludmann Marthe H.R, Andrey Y Abramov. Mitochondrial calcium imbalance in Parkinson’s disease. Neurosci Lett. 2018; 663: 86–90. PMID: 28838811 DOI: 10.1016/j.neulet.2017.08.044

19. Venediktova Natalia I., Pavlik Lyubov L, Belosludtseva Natalia V,. Khmil Natalya V, Murzaeva Svetlana V, Mironova Galina D. Formation of lamellar bodies in rat liver mitochondria in hyperthyroidism. J Bioenerg Biomembr. 2018; 50(4):289–295. PMID: 29721776 DOI: 10.1007/s10863-018-9758-8

20. Mironova G, Pavlik LL, KirovaYuI, Belosludtseva NV, Mosentsov A, Khmil NV, Germanova EL., Lukyanova LD. Effect of hypoxia on mitochondrial enzymes and ultrastructure in the brain cortex of rats with different tolerance to oxygen shortage. J Bioenerg Biomembr. 2019; 51(5):329–340.PMID: 31342235 DOI: 10.1007/s10863-019-09806-7

21. Belichenko PV, Machanov MA, Fedorov AA, Krasnov I.B., Leontovich TA. Effects of spaceflight on dendrites of the neurons of the rats brain. Physiologist. 1990 Feb; 33(1 Suppl):S12–5. PMID: 2371317

22. Belichenko PV, Leontovich TA. The giant multipolar neurons of the reticular formation in the rat brain stem after a 14-day space flight. Aviakosm Ekolog Med. 1992; 26 (5-6):24–27. PMID: 1307031.

23. Dyachkova, L.N. Ultrastructural changes in somatosensory cortex of albino rats during space flight. Biol Bull Russ Acad Sci. 2007; 34: 307–309. https://doi.org/10.1134/S106235900703015

24. Santalova Irina M, Gordon Rita Ya, Mikheeva Irina B, Khutsian Sergei S, Maevsky Eugene I. Pecularities of the structure of glycogen as an indicator of the functional state of mauthner neurons in fish Percсottus glehni during wintering. Neurosci Lett. 2018; 664:133–138. PMID: 29129679 DOI: 10.1016/j.neulet.2017.11.024

25. European Convention for the Protection of Vertebrate Animals Used for Experimental and Other Scientific Purposes. Strasbourg, 18.III.1986.

26. Behnke BJ, Stabley JN, McCullough DJ et al. Effects of spaceflight and ground recovery on mesenteric altery and vein constrictor properties in mice. FASEB J. 2013; 27(1): 339–409.

27. The ALLEN Mouse Brain Atlas (http://mouse.brain-map.org/static/atlas).

28. Fiala JC, Harris KM. Extending unbiased stereology of brain ultrastructure to three-dimensional volumes. J AmMed Inform Assoc. 2001; 8(1):1–16.PMID: 11141509 PMCID: PMC134588 DOI: 10.1136/jamia.2001.0080001

29. Moshkov DA, Mikhailova GZ, Grigorieva EE, Shtanchaev RS. Role of different dendrites in the functional activity of the central neuron controlling goldfish behavior. J Integr Neurosci. 2009; 8 (4): 441–451. PMID: 20205297 DOI: 10.1142/s0219635209002307

30. Moshkov DA, Shtanchaev RS, Mikheeva IB, Bezgina EN, Kokanova NA, Mikhailova GZ, Tiras NR, Pavlik LL. J Integr Neurosci. 2013; 12 (1):17–34. PMID: 23621454 DOI: 10.1142/S0219635213500039

